## Supplementary figures and tables for "The environmentally-regulated interplay between local three-dimensional chromatin organisation and transcription of *proVWX* in *E. coli*"

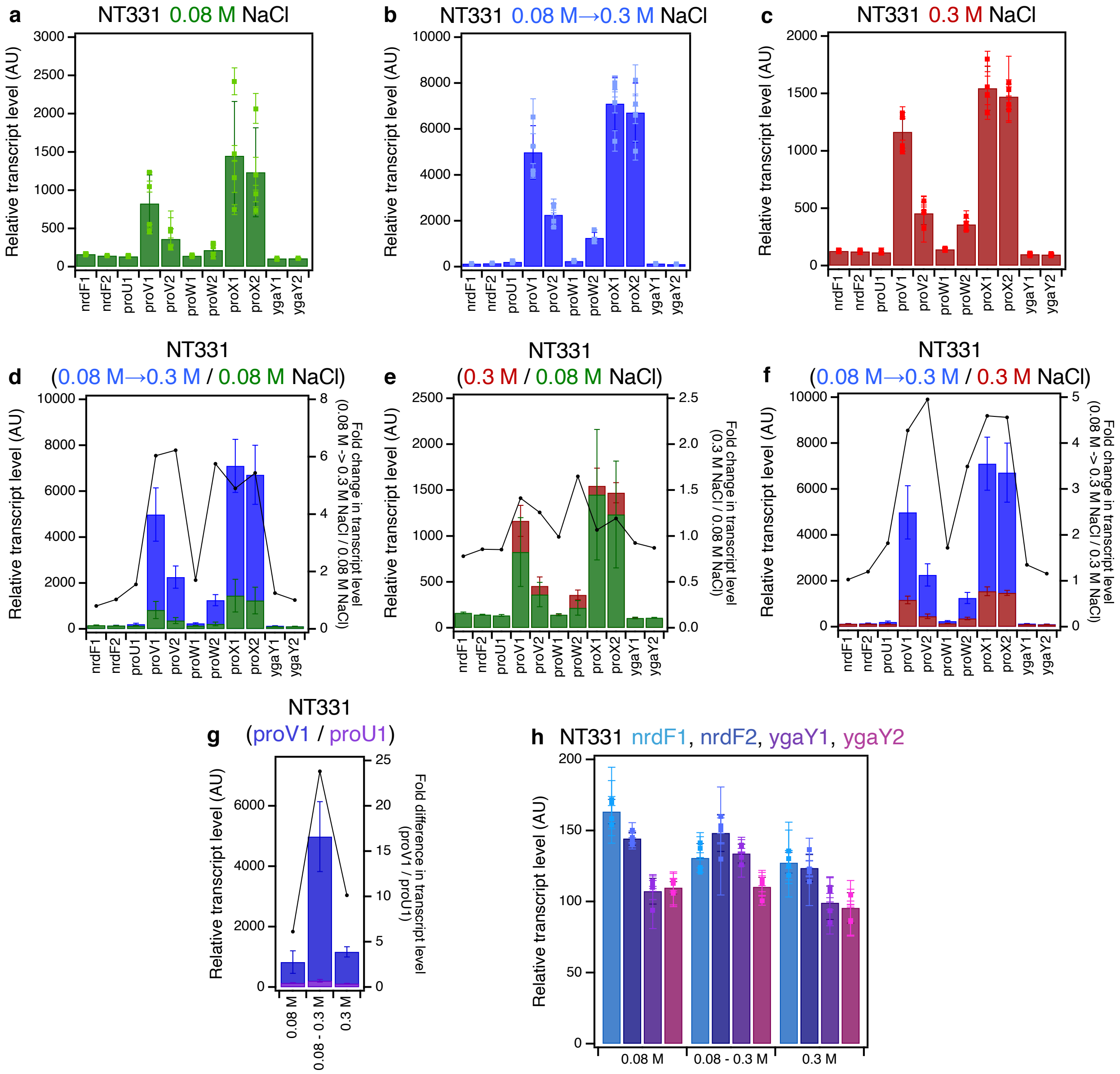


**Figure S1: The RT-qPCR profile of *proVWX* and its flanking regions in NT331** during **(a)** exponential growth in M9 medium with 0.08 M NaCl, **(b)** hyperosmotic shock in M9 medium from 0.08 M to 0.3 M NaCl, and **(c)** exponential growth in M9 medium with 0.3 M NaCl. **The fold changes in transcript levels of *proVWX* and its flanking regions** between **(d)** a hyperosmotic shock and exponential growth at 0.08 M NaCl, **(e)** exponential growth at 0.3 M NaCl and 0.08 M NaCl, and **(f)** a hyperosmotic shock and exponential growth at 0.3 M NaCl. **(g) The difference in the transcript level of the *proV*1 amplicon compared to the *proU*1 amplicon** during exponential growth at 0.08 M NaCl, following a hyperosmotic shock, and during exponential growth at 0.3 M NaCl. **(h) The transcript levels of amplicons flanking *proVWX*** during exponential growth at 0.08 M NaCl, following a hyperosmotic shock, and during exponential growth at 0.3 M NaCl. **Y-axes:** All bar graphs and data points with error bars show relative transcript levels in arbitrary units and are plotted on the left y-axis. Plots without error bars show fold changes in transcript levels and correspond to the right y-axis. **Internal control: *hcaT*.** See also Figure 1. **Error bars represent standard deviation. n=3 for dot plots, n=4 for bar graphs.**

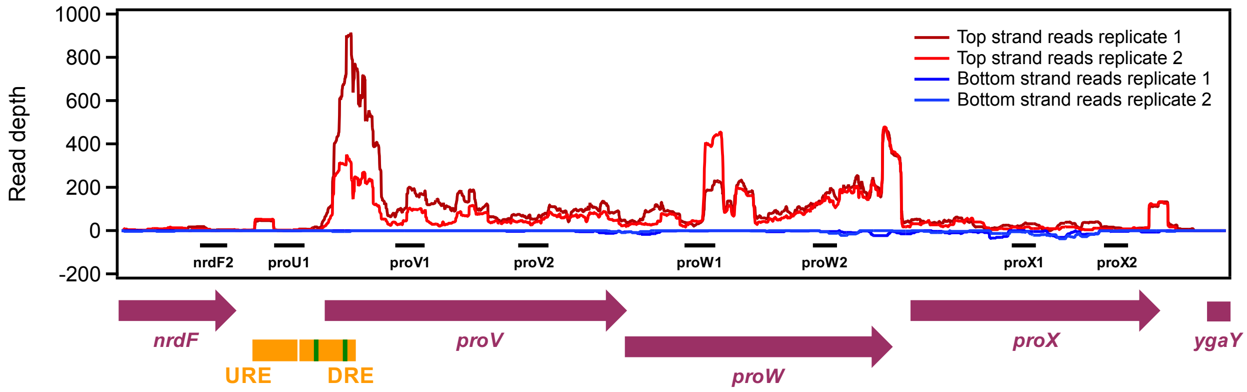


**Figure S2: The Term-seq profile of the *proVWX* operon shows the presence of transcription termination sites downstream of *proW*1, and between *proW*2 and *proX*1.**

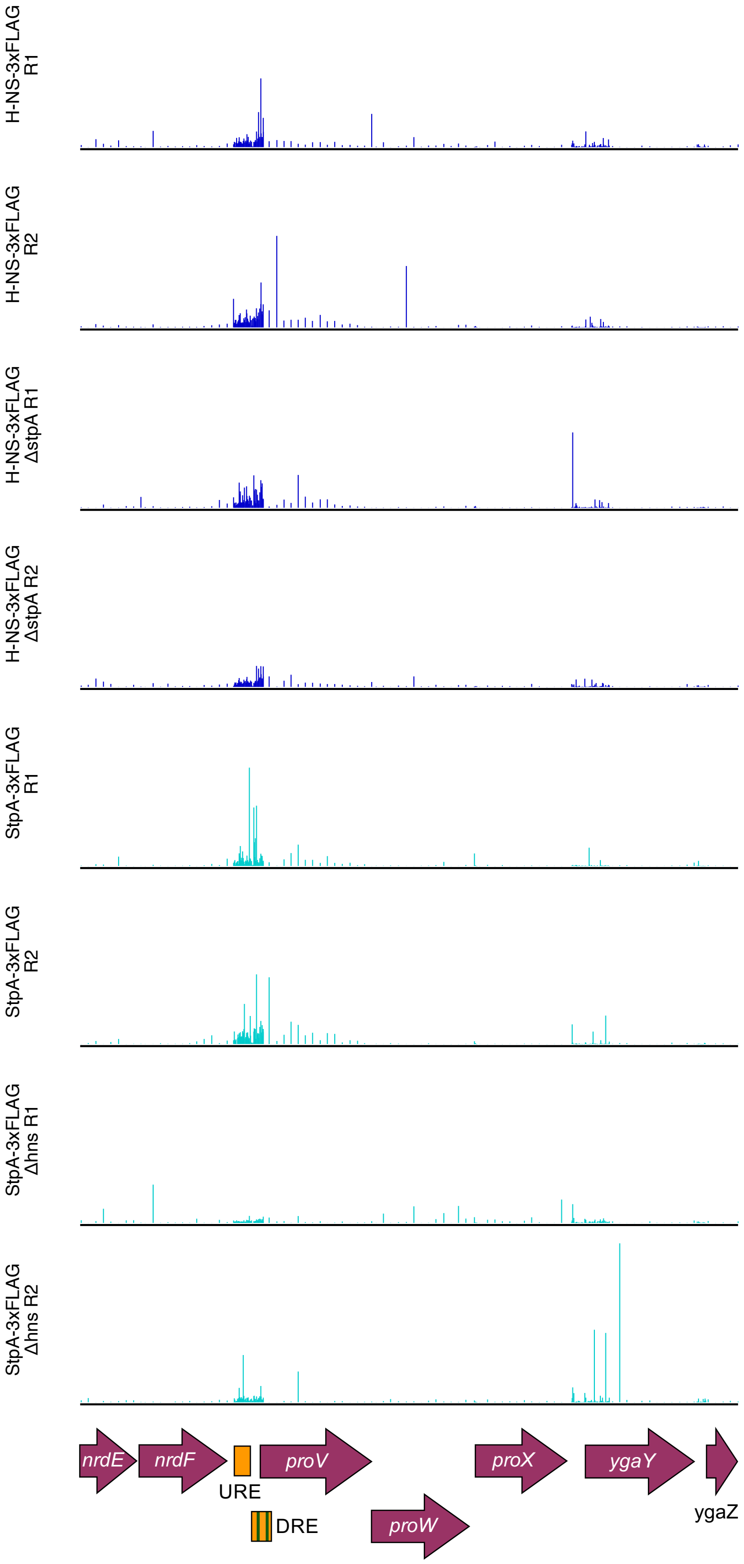


**Figure S3: The chromatin immunoprecipitation (ChIP) profiles of H-NS-3xFLAG and StpA-3XFLAG at *proVWX*[1]. H-NS-3xFLAG:** H-NS occupies the regulatory region of *proVWX* and the *proV* ORF. H-NS also binds the cryptic *ygaY* gene positioned downstream of *proVWX*. **H-NS-3xFLAG Δ*stpA*:** The occupancy and distribution of H-NS on *proVWX* is not affected by the absence of StpA within the H-NS—DNA nucleoprotein structure. **StpA-3xFLAG:** The occupancy of StpA on *proVWX* overlaps with the distribution of H-NS. The H-NS—DNA nucleoprotein at *proVWX* is interspersed with StpA. **StpA-3xFLAG Δ*hns*:** The distribution of StpA on *proVWX* is affected by the absence of H-NS. In a Δ*hns* background, StpA occupies the regulatory elements upstream of the *proV* ORF, but the nucleoprotein structure on the *proV* ORF is lost. **R1 and R2** are biological replicates. The purple arrows represent ORFs. The orange bars mark the *proVWX* upstream and downstream regulatory elements (URE and DRE). The green bars within the DRE designate high-affinity H-NS binding sites.

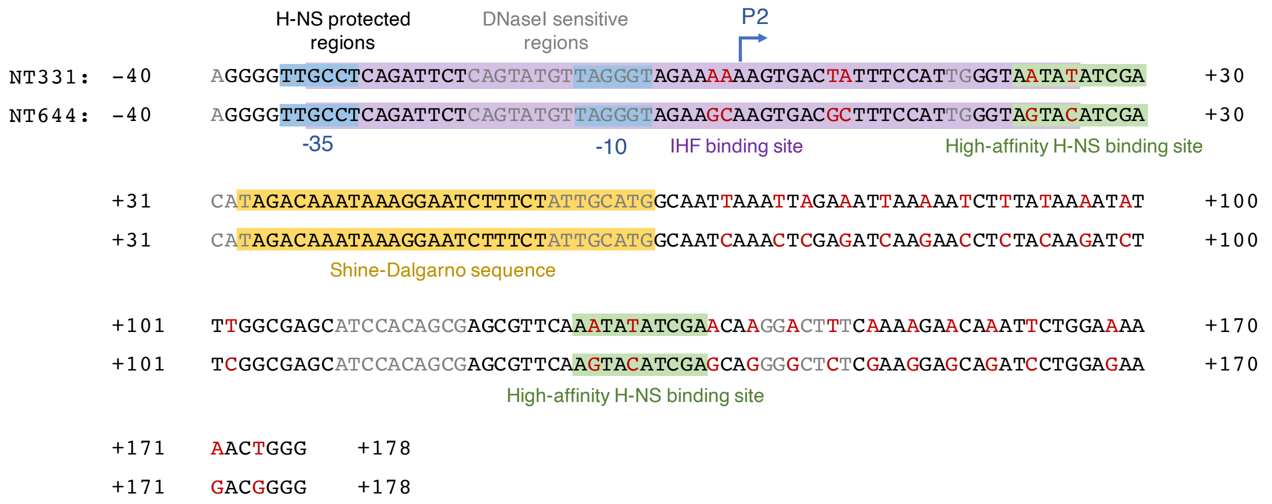


**Figure S4: A comparison between the wild-type *proVWX* DRE in NT331 and the mutant DRE in NT644.** H-NS protected and DNaseI sensitive regions of the DRE as determined in REF[2] are represented in black and grey, respectively. Point mutations to the DRE are marked in red. The P2 -35 sequence, -10 sequence, and TSS are in blue. The Shine-Dalgarno sequence is coloured yellow. The pair of high-affinity H-NS binding sites of the DRE[3] are highlighted in green, and the IHF binding site encompassing P2 is in purple.

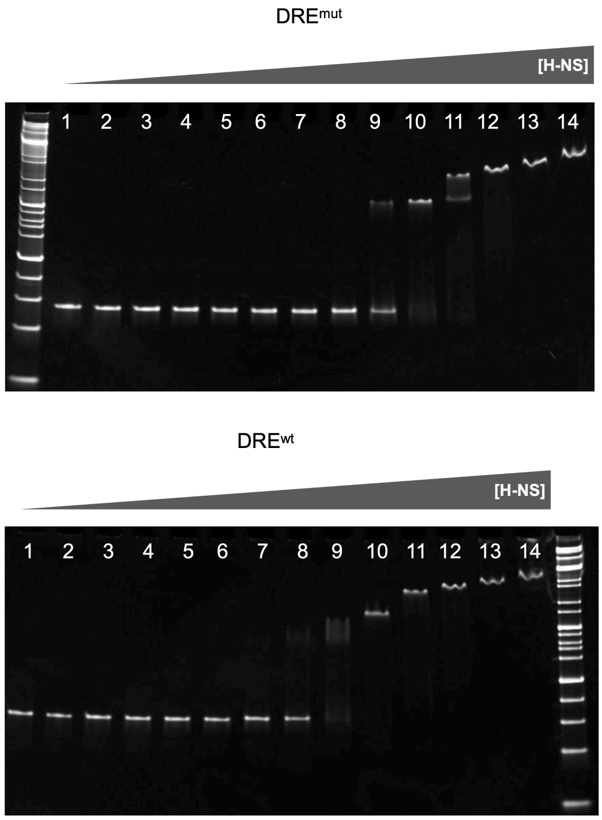


| Lane | [H-NS] (nM) |
|---|---|
| 1 | 31.7 |
| 2 | 52.8 |
| 3 | 88.0 |
| 4 | 146.6 |
| 5 | 244.4 |
| 6 | 407.3 |
| 7 | 678.8 |
| 8 | 1131 |
| 9 | 1885 |
| 10 | 3142 |
| 11 | 5238 |
| 12 | 8730 |
| 13 | 14550 |
| 14 | 24250 |

**Figure S5: The mutated DRE (top) has a lower affinity for H-NS than the wild-type DRE (bottom)**

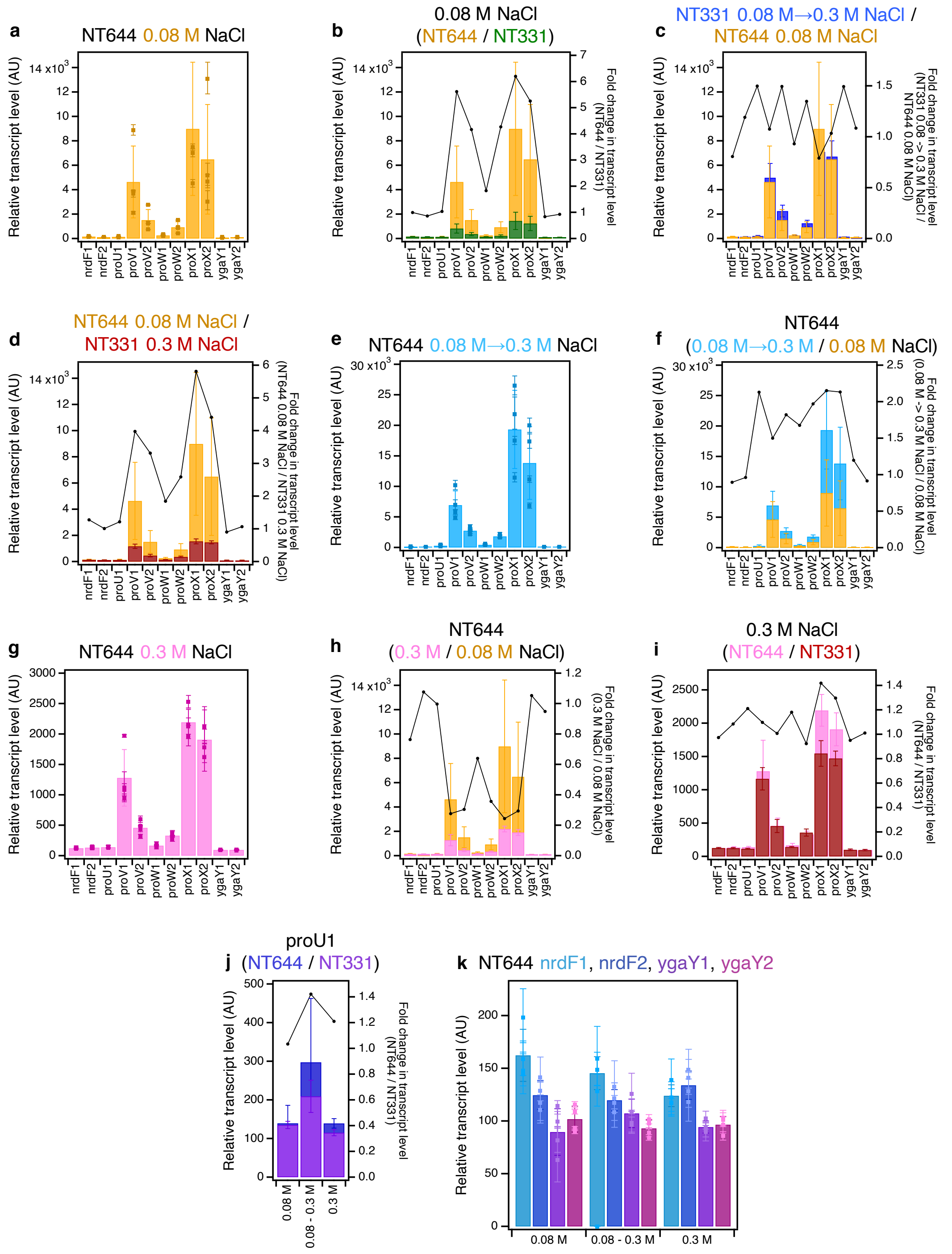

**Figure S6: The RT-qPCR profile of *proVWX* and its flanking regions in NT644 (a)** during exponential growth in M9 medium with 0.08 M NaCl. **A comparison of transcript levels of *proVWX* and its flanking regions (b)** between NT644 and NT331 during exponential growth in M9 medium with 0.08 M NaCl, **(c)** between NT644 growing exponentially at 0.08 M NaCl and NT331 subjected to a hyperosmotic shock from 0.08 M to 0.3 M NaCl, and **(d)** between NT644 at 0.08 M NaCl and NT331 at 0.3 M NaCl. **The RT-qPCR profile of *proVWX* and its flanking regions in NT644 (e)** after a hyperosmotic shock from 0.08 M to 0.3 M NaCl, and **(f)** the fold change in transcript levels compared to exponential growth at 0.08 M NaCl. **The RT-qPCR profile of *proVWX* and its flanking regions (g)** during exponential growth in M9 medium with 0.3 M NaCl, and a comparison of this profile with that of **(h)** exponential growth of NT644 in M9 medium with 0.08 M NaCl, and **(i)** exponential growth of NT331 in M9 medium with 0.3 M NaCl. **(j) The fold change in transcript level of the *proU*1 amplicon** between NT644 and NT331 during exponential growth at 0.08 M NaCl, following a hyperosmotic shock, and during exponential growth at 0.3 M NaCl. **(k) The relative transcript level in NT644 at amplicons flanking *proVWX*** during exponential growth at 0.08 M NaCl, following a hyperosmotic shock, and during exponential growth at 0.3 M NaCl. **Y-axes:** All bar graphs and data points with error bars show relative expression levels in arbitrary units and are plotted on the left y-axis. Plots without error bars show fold-change in expression level and correspond to the right y-axis. **Internal control: *hcaT*.** See also Figure 2. **Error bars represent standard deviation. n=3 for dot plots, n=4 for bar graphs.**

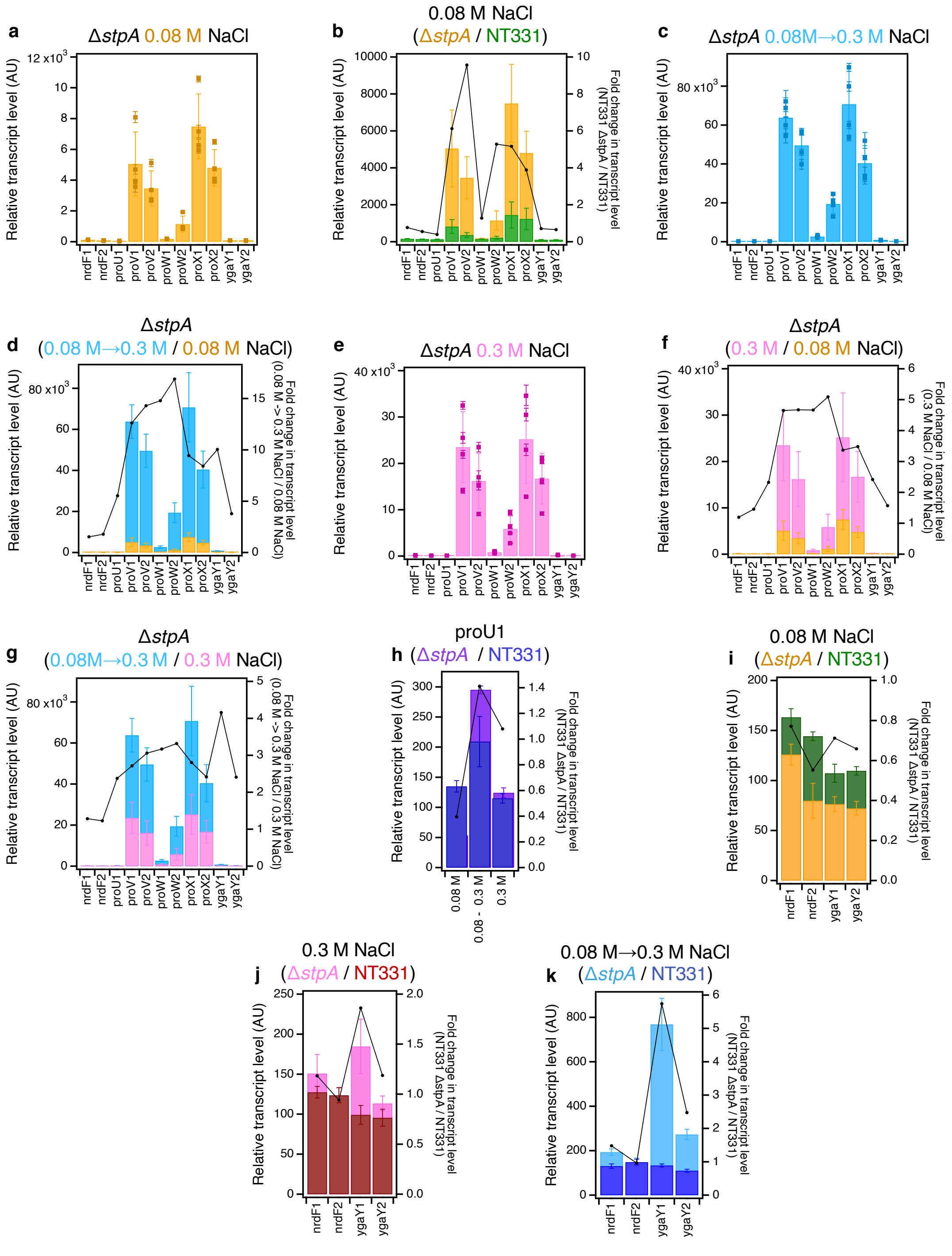


**Figure S7: The RT-qPCR profile of the *proVWX* operon and its flanking regions in NT331 Δ*stpA* (a)** during exponential growth at 0.08 M NaCl and **(b)** the fold-change in the transcript levels of the amplicons compared to NT331. **(c) The RT-qPCR profile of the *proVWX* operon and its flanking regions in NT331 Δ*stpA*** upon a hyperosmotic shock from 0.08 M to 0.3 M NaCl, and **(d)** the fold-change in transcript levels of the amplicons in comparison to exponential growth at 0.08 M NaCl. **(e) The RT-qPCR profile of the *proVWX* operon and its flanking regions in NT331 Δ*stpA*** during exponential growth at 0.3 M NaCl, and the fold-change in transcript levels of the amplicons with respect to **(f)** exponential growth at 0.08 M NaCl, and **(g)** a hyperosmotic shock. **(h) The fold difference in transcript level of amplicon *proU*1** between NT331 Δ*stpA* and NT331. **The fold change in transcript levels of the *nrdF* and *ygaY* amplicons between NT331 Δ*stpA* and NT331 (i)** during exponential growth at 0.08 M NaCl, **(j)** exponential growth at 0.3 M NaCl, and **(k)** following a hyperosmotic shock. **Y-axes:** All bar graphs and data points with error bars show relative transcript levels in arbitrary units and are plotted on the left y-axis. Plots without error bars show fold-change in transcript level and correspond to the right y-axis. **Internal control: *hcaT*.** See also Figure 3. **Error bars represent standard deviation. n=3 for dot plots, n=4 for bar graphs.**

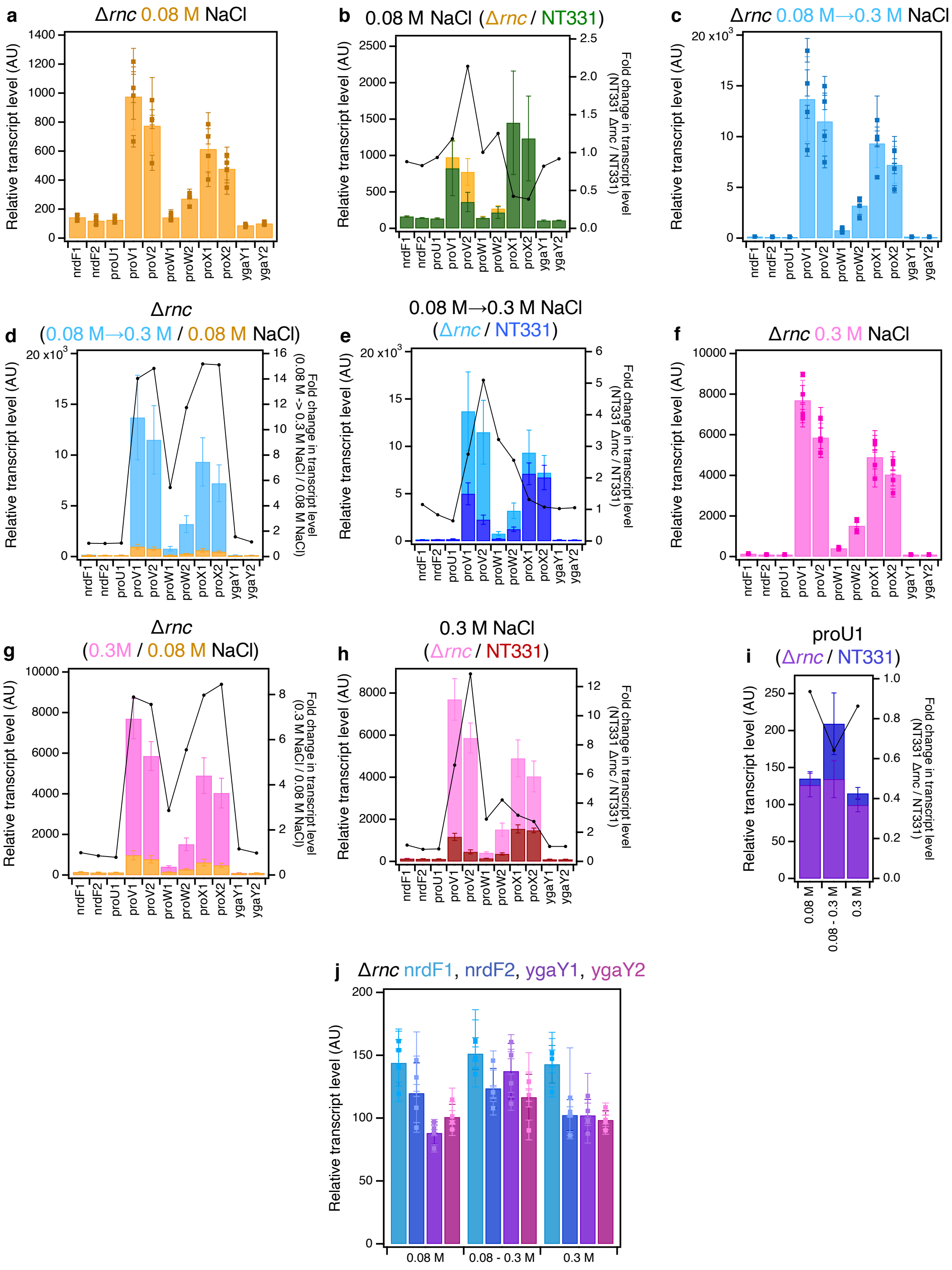


**Figure S8: The RT-qPCR profile of *proVWX* and its flanking regions in NT331 Δ*rnc*** during **(a)** exponential growth at 0.08 M NaCl, and the fold change in transcript levels of the amplicons **(b)** compared to NT331 growing exponentially at 0.08 M NaCl. **The RT-qPCR profile of *proVWX* and its flanking regions in NT331 Δ*rnc*** upon **(c)** a hyperosmotic shock from 0.08 M NaCl to 0.3 M NaCl, and the fold change in transcript levels of the amplicons in comparison to **(d)** NT331 Δ*rnc* growing exponentially at 0.08 M NaCl, and **(e)** NT331 following a hyperosmotic shock. **The RT-qPCR profile of *proVWX* and its flanking regions in NT331 Δ*rnc*** during **(f)** exponential growth at 0.3 M NaCl, and the fold change in transcript leve;s of the amplicons relative to **(g)** NT331 Δ*rnc* growing exponentially at 0.08 M NaCl, and **(h)** NT331 growing exponentially at 0.3 M NaCl. **(i) The fold change in the relative transcript levels of the *proU*1 amplicon** between NT331 Δ*rnc* and NT331. **(j) The relative transcript level in NT331 Δ*rnc* at amplicons flanking *proVWX*** during exponential growth at 0.08 M NaCl, following a hyperosmotic shock, and during exponential growth at 0.3 M NaCl. **Y-axes:** All bar graphs and data points with error bars show relative transcript levels in arbitrary units and are plotted on the left y-axis. Plots without error bars show fold-change in transcript level and correspond to the right y-axis. **Internal control: *hcaT*.** See also Figure 4. **Error bars represent standard deviation. n=3 for dot plots, n=4 for bar graphs.**

**_Escherichia coli_ cells show global differences in the chromosome contact profiles during growth at different osmolarity conditions.**

The binding of NAPs to DNA is sensitive to environmental conditions such as pH, temperature, and osmolarity [19–28]. Consequently, changes to the ambient growth conditions of bacteria are reflected in an altered NAP binding profile of the chromosome, and hence, in the three-dimensional chromosome organization. We first used Hi-C, a high-throughput chromosome conformation capture technique, to examine the global differences in the chromosome contact profiles of MG1655 Δ*endA* (NT331) (Figures S10 and S11) during growth in a low-salt (0.08 M NaCl) medium, following a hyperosmotic shock (0.08 M -> 0.3 M NaCl), and in a high-salt (0.3 M NaCl) medium (Figure S9). To improve the signal-to-noise ratio of the chromosome contact maps, *E. coli* cells were permeabilised with methanol prior to formaldehyde fixation, and the proximity ligation step of Hi-C was performed using the insoluble fraction of digested, cross-linked chromatin (Figure S14 and S15) [29].

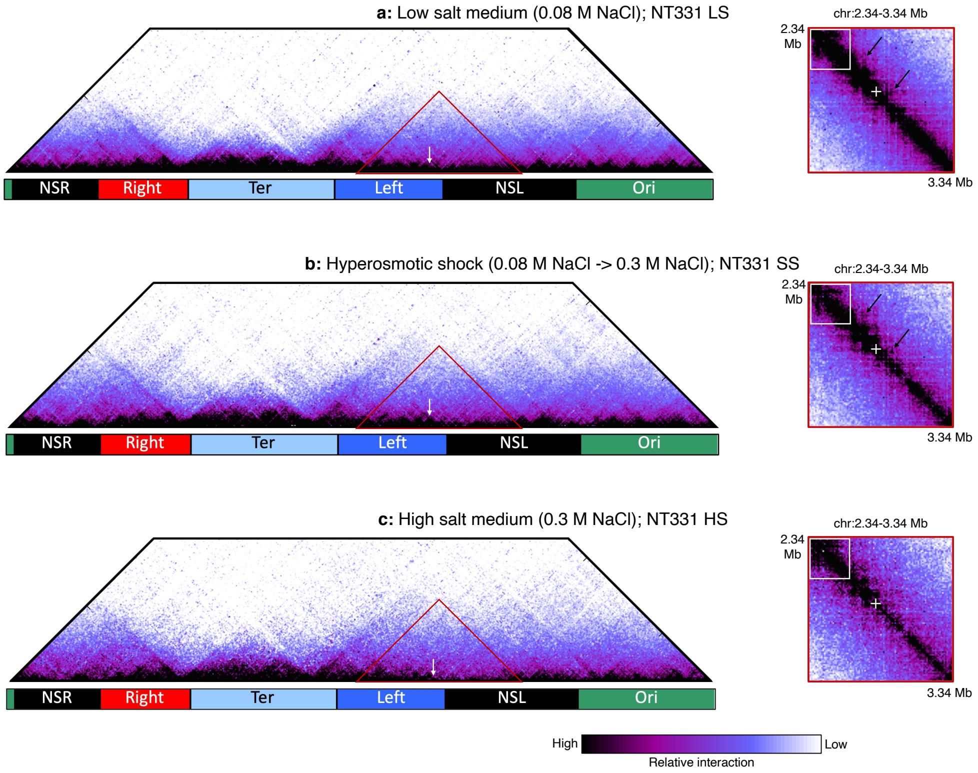


**Figure S9: _Escherichia coli_ cells show global differences in chromosome contact profiles during growth at different osmolarity conditions. (a) & (b)** The chromosome around *proVWX* (marked with a red triangle in the left panels, shown in the right panels) decompacts locally when *E. coli* cells growing exponentially in a low salt medium **(a)** are subjected to a hyperosmotic shock **(b)**. The position of *proVWX* is marked with white arrows in the left panels and with white '+' marks in the right panels. The local chromosome maintains features of the finer chromosome organization after a hyperosmotic shock, such as loops (black arrows, right panels) and the arrow-like structure at 2.37 Mb (white squares, right panels). **(c)** Upon adaptation to hyperosmotic stress represented as exponential growth in a high salt medium, loci either show decompaction compared to growth in a low salt medium such as the chromatin encompassing, and positioned locally downstream of *proVWX*, or show a stronger compaction, for example, the region encompassing the arrow-like structure at 2.37 Mb (white square, right panels). **Organism:** *Escherichia coli* MG1655 Δ*endA* (NT331); **3C-based study:** Hi-C; **Resolution:** 10 kb; **Fixation conditions:** 80% cold methanol for 10 minutes followed by 3% formaldehyde for 1 hour (Figures S14-S15); **Restriction enzyme:** PsuI (ThermoFisher Scientific); **Fractionation:** Yes.

The NT331 chromosome contact maps show global chromosomal rearrangements in response to osmolarity. The rearrangements are also observed in the vicinity of the osmosensitive *proVWX* operon (Figure S9). When *E. coli* cells in a low-salt medium are subjected to a hyperosmotic shock, the local chromosome at *proVWX* decompacts while maintaining features of the finer chromosome organization, such as loops (marked with black arrows, Figures S9a-S9b, right panels) and the arrow-like structure at 2.37 Mb (marked with white squares, Figures S9a-S9b, right panels). In a high salt environment – a condition that reflects the adaptation of *E. coli* to higher osmolarity following a hyperosmotic shock – loci either decompact further such as the chromatin encompassing, and positioned locally downstream of *proVWX*, or show a stronger compaction compared to growth in a low-salt medium, for instance, the region encompassing the arrow-like structure at 2.37 Mb (marked with a white square, Figure S9c, right panel).

**Degradation of *Escherichia coli* chromatin during 3C-based library preparation is overcome by the deletion of *endA*.**

The *Escherichia coli* K-12 MG1655 strain closely resembles the genetic make-up of archetype *E. coli* and is used as a reference for genome-wide studies of NAP-binding profiles and transcription. The strain is, therefore, the optimal choice to study the interplay between three-dimensional chromatin organisation, NAP distribution, and gene expression. However, chromatin extracted from MG1655 underwent considerable degradation during the initial steps of 3-C and Hi-C library preparation. The degradation was not observed during the lysis and solubilisation steps [4] that were carried out in a buffer with 1.0 mM EDTA but occurred extensively once the cell lysate was diluted in a restriction digestion mix with a final EDTA concentration of 0.1 mM (Figure S10a). The dependence of chromatin degradation on the concentration of EDTA, and hence, the availability of divalent ions implied that the degradation was enzymatic.

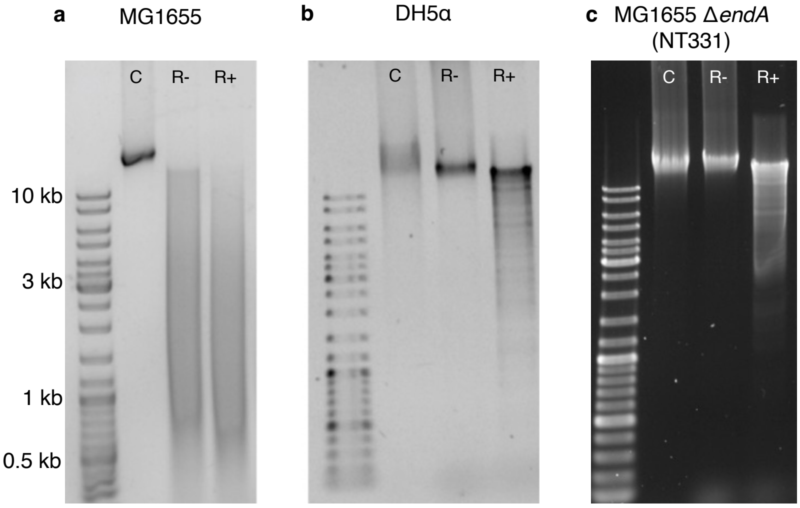


**Figure S10: Degradation of *Escherichia coli* chromatin during 3C-based library preparation is overcome by *endA* deletion. Lane C:** Chromatin preparation in 1X TE (EDTA concentration: 1.0 mM); **Lane R-:** Chromatin preparation after a 3-hour incubation in 1X restriction digestion buffer (EDTA concentration: 0.1 mM); **Lane R+:** Chromatin preparation after a 3-hour treatment with 0.4 U/µL of BglII in 1X restriction digestion buffer (EDTA concentration: 0.1 mM). **(a) MG1655** chromatin undergoes extensive degradation in a restriction digestion buffer with a final EDTA concentration of 0.1 mM. The degradation is observed as a smear in lanes R- and R+. Similar degradation is not observed in *endA* knock-out strains **(b) DH5α** and **(c) NT331**, where extracted chromatin (lane C) still runs as a heavy >10 kb band after a 3-hour incubation in a buffer with 0.1 mM EDTA (lane R-). Fixed chromatin extracted from the *endA*$^-$ strains can be digested by restriction enzymes (shown: BglII).

Endonuclease-I is a DNA-specific nuclease localised in the periplasm [5] that digests dsDNA in a sequence independent manner. It is responsible for the low quality of plasmid DNA preparations from *endA*$^+$ *E. coli* strains [6,7]. To investigate whether the enzyme also contributes to the degradation of chromatin in lysates of formaldehyde-treated cells, the stability of fixed chromatin extracted from DH5α – an *endA*$^-$ strain of *E. coli* [7] – during the initial steps of chromosome conformation capture was tested. Agarose gel electrophoresis showed that DH5α chromatin preparations do not degrade in the restriction digestion buffer with 0.1 mM EDTA (Figure S10b). Attempts to thermally denature endonuclease-I and overcome chromatin degradation were not pursued extensively since the conditions that reliably decreased degradation also promote reverse cross-linking of the chromatin and thereby interfere with proximity ligation in later steps of 3C-based protocols (Figure S11).

Therefore, MG1655 Δ*endA* (henceforth referred to as NT331) was generated using the λ-red recombinase mediated gene replacement strategy [8,9]. Chromatin preparations from fixed NT331 do not degrade when incubated in a buffer with a low concentration of EDTA (Figure S10c). Thus, all 3C-based experiments, and the associated RT-qPCR studies, were carried out in a Δ*endA* background.

Using higher concentrations of formaldehyde for fixation, for instance, 7%, also reduces chromatin degradation [10,11]. Nevertheless, we preferred to use *endA-* strains for our 3C-based studies since this reduces the potential for introducing artefacts that are associated with using high concentrations of fixatives. It also provides a wider window to finetune fixation conditions.

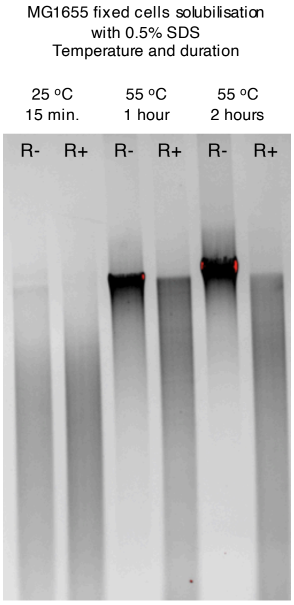


**Figure S11: Cell solubilisation conditions that reliably decrease degradation of MG1655 chromatin in low [EDTA] buffers also promote de-crosslinking of formaldehyde-fixed chromatin.** Raising the temperature and increasing the duration of 0.5% SDS treatment during lysis and solubilisation of fixed cells increases the stability of the extracted chromatin in low [EDTA] buffer.

**RNA preparations do not show detectable genomic DNA contamination.**

In compliance with MIQE guidelines [12], all RNA preparations were tested for genomic DNA contamination. ~100 ng of RNA with and without RNase treatment (RNase+ and RNase-, respectively) were visualised on a 1.2% agarose gel pre-stained with 1X GelRed (Sigma-Aldrich). The absence of a nucleic acid signal in the RNase+ wells shows the absence of detectable genomic DNA contamination (Figure S12).

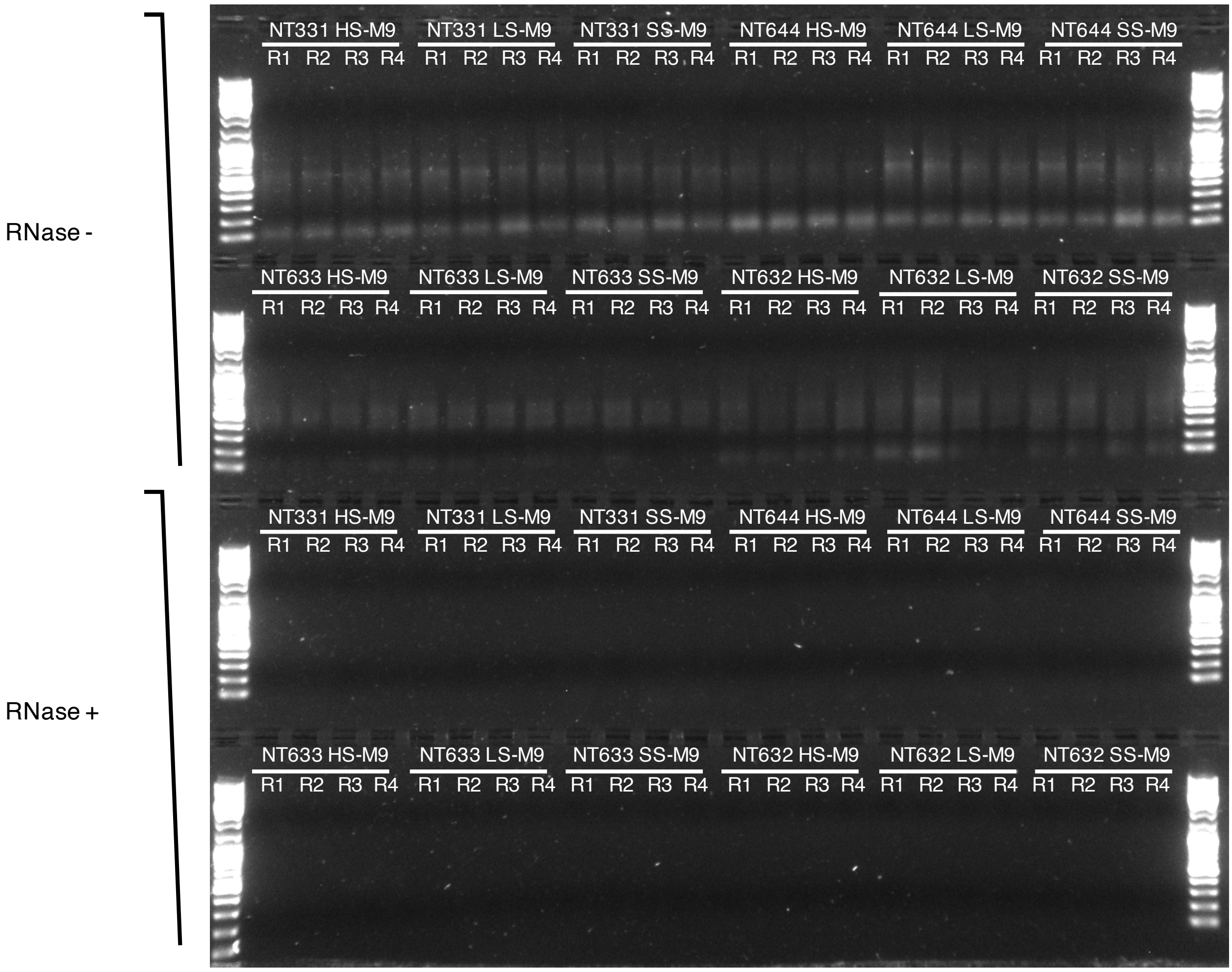


**Figure S12: DNA contamination assessment of RNA preparations.** RNA preparations of NT331, NT644, NT633, and NT632 (Table 1) show no nucleic acid signal upon treatment with RNase.

**The specificity of primer pairs used for RT-qPCR studies was determined with melting curve analysis and Sanger sequencing of the amplified product.**

In compliance with MIQE guidelines [12], the specificity of primer pairs was experimentally determined by Sanger sequencing (BaseClear B.V., Leiden, The Netherlands) of the amplified products (SI 1B and SI 2). The melting curve profiles and the melting temperature (Tm) of the sequenced amplicons (Figure S13) were used to gauge the specificity of amplification in RT-qPCR experiments.

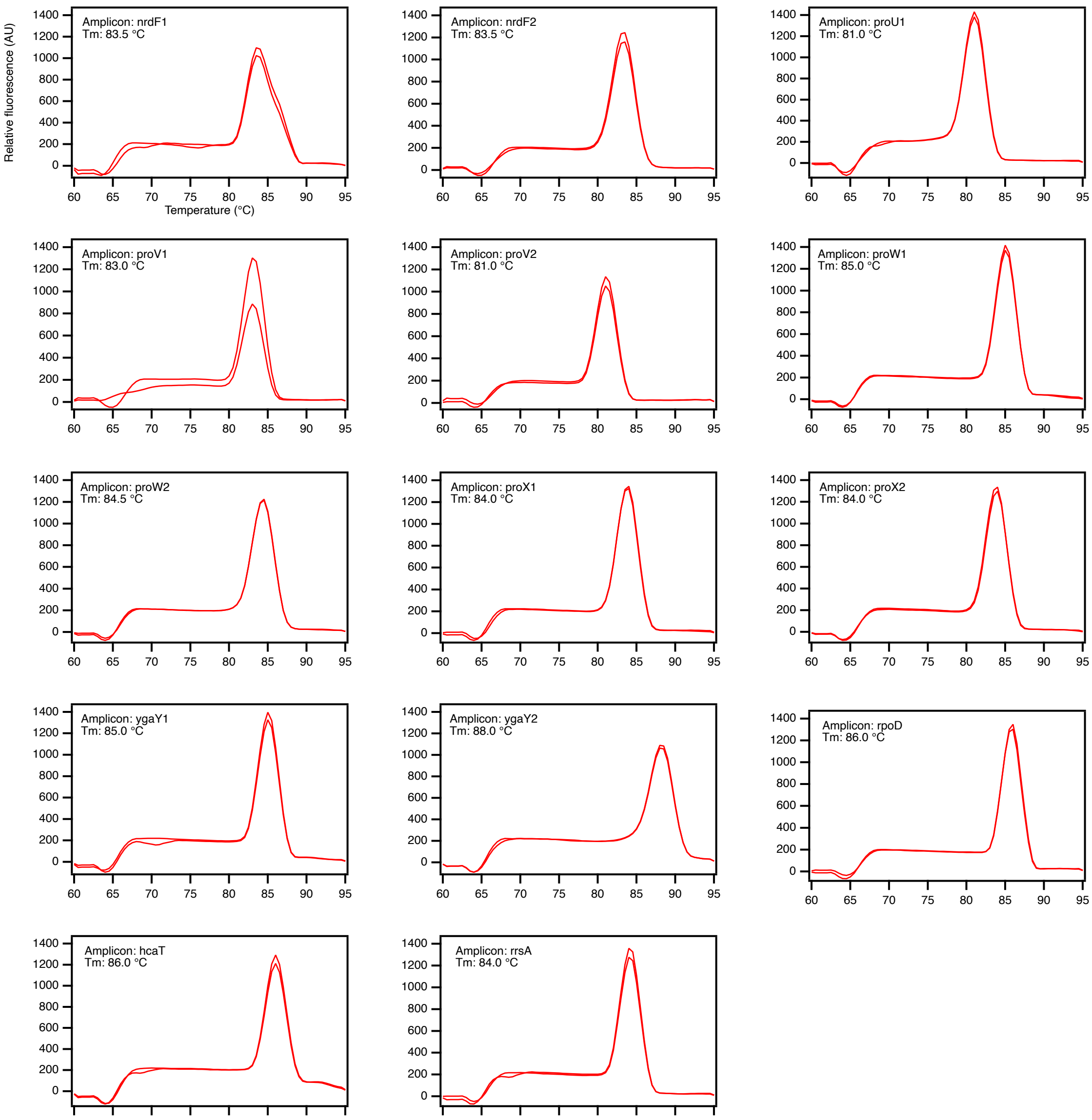


**Figure S13: RT-qPCR amplicon melting curves.** The specificity of amplification in RT-qPCR reactions was determined from the melting curve of the fragment amplified in each well. The melting curve and the melting temperature (Tm) of the amplicons reported on in this manuscript are shown here. The sequences of the fragments used for this experiment were verified with Sanger sequencing (SI 1B and SI 2).

**Proximity ligation with the insoluble fraction of digested, cross-linked chromatin, and methanol permeabilization of *Escherichia coli* prior to formaldehyde fixation improves the signal-to-noise ratio in chromosome contact maps.**

Hi-C libraries of NT331 fixed with 3% formaldehyde have a low signal-to-noise ratio (Figure S14a), indicating inefficient formaldehyde-mediated cross-linking. Taking an earlier report of 3C-based studies in *E. coli* [10] into account, we raised the concentration of formaldehyde for fixation from 3% to 7%. However, the change did not contribute to a significant improvement in chromosome contact maps (Figure S15).

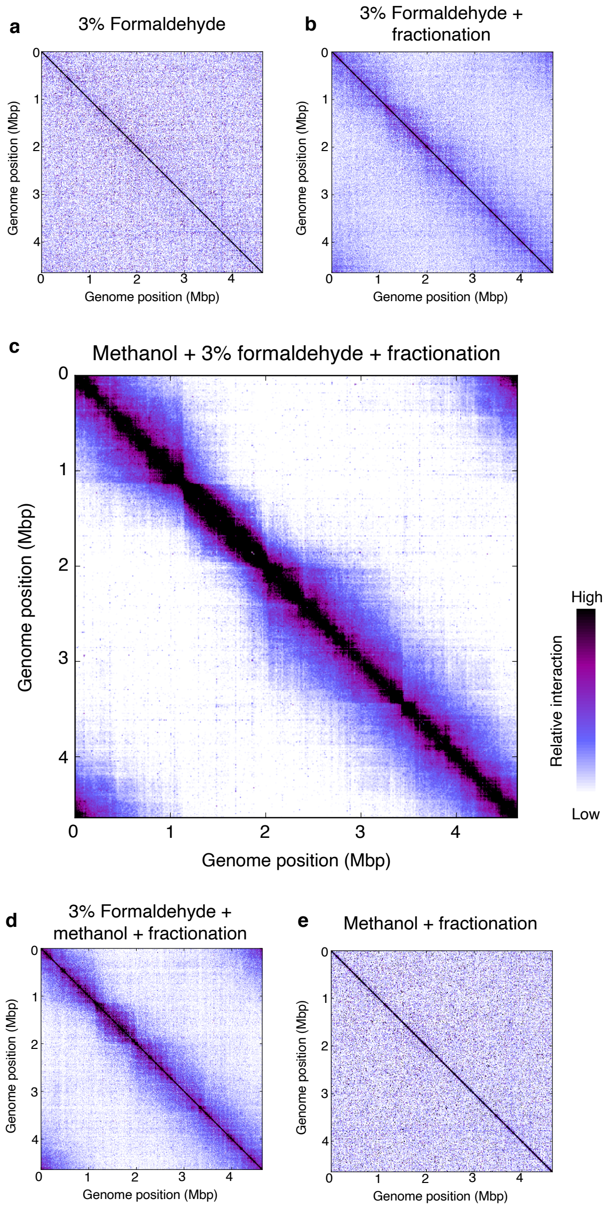


**Figure S14: Proximity ligation with the insoluble fraction of digested, cross-linked chromatin, and methanol permeabilization of *Escherichia coli* prior to formaldehyde fixation improves the signal-to-noise ratio in chromosome contact maps. (a)** Hi-C libraries prepared from cells fixed with 3% formaldehyde have a low signal-to-noise ratio. **(b)** The signal-to-noise ratio is improved by proximity ligation with the insoluble fraction of digested, cross-linked chromatin and **(c)** further improved by methanol permeabilization of *E. coli* cells prior to formaldehyde fixation**. (d)** Only a marginal improvement in the signal-to-noise ratio of chromosome contact maps is observed when methanol treatment is performed after formaldehyde fixation. **(e)** *E. coli* cells permeabilised with methanol but not fixed with formaldehyde cannot be used to map chromosome structure. **Organism:** *Escherichia coli* MG1655 Δ*endA* (NT331); **3C-based study:** Hi-C; **Resolution:** 10 kb; **Growth conditions:** LB medium, 37 $^{o}$C, exponential phase; **Fixation conditions: a and b:** 3% formaldehyde, 1 hour, **c:** 80% cold methanol for 10 minutes followed by 3% formaldehyde for 1 hour, **d:** 3% formaldehyde for 1 hour followed by 80% cold methanol for 10 minutes, **e:** 80% cold methanol for 10 minutes; **Restriction enzyme:** PsuI (ThermoFisher Scientific); **Fractionation: a:** No, **b-e:** Yes.

A low signal-to-noise ratio in chromosome contact maps may arise from ligation between freely moving, non-crosslinked DNA molecules [13]. This effect can be overcome by fractionating the digested, cross-linked chromatin into its supernatant, and pellet fractions by centrifugation [13,14]. Cross-linked DNA-protein complexes are enriched in the pellet and freely moving DNA molecules in the supernatant. This allows contact maps with a high signal-to-noise ratio to be generated when proximity ligation is carried out with only the pellet fraction [13,14]. In agreement with previous observations in E14.5 mouse embryos [14], *Saccharomyces cerevisiae* [13], and *S. pombe* [13], incorporating fractionation and using only the pellet fraction for proximity ligation improved the signal-to-noise ratio of the *Escherichia coli* NT331 contact map (Figure S14b). Fractionation was also incorporated in 3C-based studies of *Bacillus subtilis* [15].

Formaldehyde is a standard fixative in histology and (immuno-)histochemical studies where it is used either alone, or in combination with methanol [16,17]. Methanol dissolves lipids from cell membranes and coagulates proteins, thus, simultaneously permeabilizing and fixing histological preparations [17,18]. We extrapolated this to *E. coli*, and permeabilized the cells with 80% methanol prior to formaldehyde fixation. This treatment led to a significant improvement in the signal-to-noise ratio of chromosome contact maps (Figure S14c). Contact maps of chromosomes fixed in this manner are qualitatively indistinguishable from those fixed with 7% formaldehyde as in REF[10]. In comparison, only a weak improvement in the signal-to-noise ratio of chromosome contact maps was observed when methanol treatment was performed after formaldehyde fixation (Figure S14d). Methanol treatment alone could not be used to study chromosome conformation (Figure S14e).

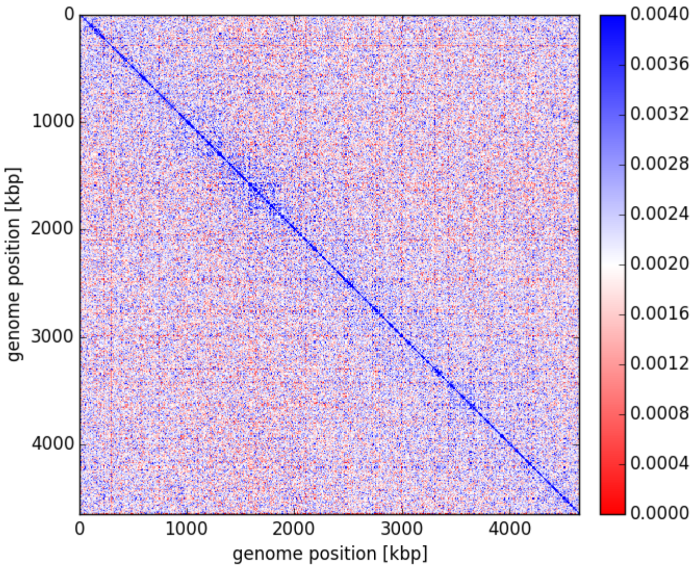


**Figure S15: Fixation of *E. coli* cells with higher concentrations of formaldehyde does not significantly improve the signal-to-noise ratio in chromosome contact maps.** Chromosome contact maps of *E. coli* cells fixed using 7% formaldehyde do not show a significant improvement in the signal-to-noise ratio compared to maps generated from *E. coli* cells fixed with 3% formaldehyde (Figure S14a). **Organism:** *Escherichia coli* MG1655 Δ*endA* (NT331); **3C-based study:** 3C-Seq; **Growth conditions:** LB medium, 37 °C, exponential phase; **Fixation conditions:** 7% formaldehyde, 1 hour; **Restriction enzyme:** HpaII (NEB); **Fractionation:** Not performed.

### Supplementary tables

**Table S1: Relative transcript levels of amplicons within and flanking the *proVWX* operon in NT331. Internal control: *rpoD***

| Amplicon | NT331 0.08 M NaCl | NT331 Hyperosmotic shock | NT331 0.3 M NaCl |
|---|---|---|---|
| nrdF1 | 81.76±8.58 | 80.31±6.67 | 85.26±11.58 |
| nrdF2 | 72.65±11.31 | 90.84±2.98 | 82.50±9.49 |
| proU1 | 67.60±7.90 | 128.77±27.04 | 76.72±7.43 |
| proV1 | 402.16±150.56 | 3047.84±637.44 | 773.91±89.45 |
| proV2 | 177.21±50.60 | 1376.57±238.83 | 304.88±73.74 |
| proW1 | 71.52±11.27 | 148.65±13.91 | 94.40±12.63 |
| proW2 | 106.71±31.29 | 766.58±137.55 | 237.86±27.42 |
| proX1 | 699.64±279.30 | 4365.00±726.11 | 1025.76±84.89 |
| proX2 | 598.77±226.02 | 4125.85±795.50 | 979.43±65.30 |
| ygaY1 | 53.92±8.70 | 82.41±8.39 | 65.80±4.53 |
| ygaY2 | 54.98±5.93 | 67.00±7.56 | 63.53±6.03 |

**Table S2: Relative transcript levels of amplicons within and flanking the *proVWX* operon in NT331. Internal control: *hcaT***

| Amplicon | NT331 0.08 M NaCl | NT331 Hyperosmotic shock | NT331 0.3 M NaCl |
|---|---|---|---|
| nrdF1 | 163.32±8.61 | 130.70±10.10 | 127.36±7.29 |
| nrdF2 | 144.37±4.35 | 148.28±12.99 | 123.56±9.44 |
| proU1 | 135.01±9.47 | 209.23±41.67 | 115.02±7.97 |
| proV1 | 825.06±374.20 | 4982.62±1159.38 | 1165.23±168.78 |
| proV2 | 362.27±132.65 | 2255.64±482.54 | 455.49±98.55 |
| proW1 | 142.41±11.69 | 242.24±25.71 | 141.03±7.61 |
| proW2 | 217.29±79.75 | 1250.22±241.01 | 358.28±53.15 |
| proX1 | 1449.73±710.92 | 7101.04±1153.99 | 1545.50±192.71 |
| proX2 | 1234.74±581.37 | 6713.35±1288.12 | 1471.58±109.99 |
| ygaY1 | 107.28±9.05 | 133.79±6.59 | 99.17±11.73 |
| ygaY2 | 109.74±4.21 | 110.35±6.71 | 95.51±10.66 |

**Table S3: Relative transcript levels of amplicons within and flanking the *proVWX* operon in NT644. Internal control: *rpoD***

| Amplicon | NT644 0.08 M NaCl | NT644 Hyperosmotic shock | NT644 0.3 M NaCl |
|---|---|---|---|
| nrdF1 | 88.27±5.12 | 91.17±17.10 | 72.81±4.94 |
| nrdF2 | 68.34±8.97 | 71.09±5.69 | 78.61±5.15 |
| proU1 | 74.79±15.20 | 169.18±76.33 | 81.69±3.77 |
| proV1 | 2428.61±1177.42 | 4032.33±868.78 | 734.54±175.17 |
| proV2 | 796.10±341.97 | 1618.03±183.18 | 264.97±35.89 |
| proW1 | 141.29±21.40 | 254.71±41.71 | 97.67±14.01 |
| proW2 | 496.96±194.66 | 1081.64±127.27 | 193.62±19.97 |
| proX1 | 4717.25±2145.92 | 11245.74±2835.34 | 1287.77±115.42 |
| proX2 | 3378.13±1791.64 | 7965.02±2759.61 | 1128.00±210.66 |
| ygaY1 | 48.54±9.77 | 63.73±8.35 | 55.60±5.71 |
| ygaY2 | 55.56±4.41 | 55.20±3.46 | 56.70±3.78 |

**Table S4: Relative transcript levels of amplicons within and flanking the *proVWX* operon in NT644. Internal control: *hcaT***

| Amplicon | NT644 0.08 M NaCl | NT644 Hyperosmotic shock | NT644 0.3 M NaCl |
|---|---|---|---|
| nrdF1 | 162.28±24.75 | 145.43±15.62 | 123.93±10.31 |
| nrdF2 | 124.49±13.55 | 119.73±10.06 | 133.98±13.44 |
| proU1 | 139.59±46.30 | 297.39±165.21 | 139.22±12.29 |
| proV1 | 4632.45±2942.73 | 6948.43±2328.41 | 1279.27±465.74 |
| proV2 | 1508.89±862.37 | 2748.93±522.37 | 458.92±116.22 |
| proW1 | 260.22±58.78 | 436.10±120.87 | 166.54±27.97 |
| proW2 | 927.22±448.16 | 1826.21±278.49 | 330.78±47.92 |
| proX1 | 8984.84±5455.14 | 19333.58±6401.31 | 2193.00±238.20 |
| proX2 | 6489.04±4488.92 | 13841.45±5963.81 | 1907.92±247.87 |
| ygaY1 | 89.55±22.12 | 107.22±13.53 | 94.29±5.71 |
| ygaY2 | 101.75±12.28 | 93.06±7.94 | 96.49±7.63 |

**Table S5: Relative transcript levels of amplicons within and flanking the *proVWX* operon in NT331 Δ*stpA* (NT633). Internal control: *rpoD***

| Amplicon | NT633 0.08 M NaCl | NT633 Hyperosmotic shock | NT633 0.3 M NaCl |
|---|---|---|---|
| nrdF1 | 46.25±2.53 | 79.10±7.59 | 54.52±11.33 |
| nrdF2 | 29.79±8.16 | 58.43±11.39 | 43.00±13.54 |
| proU1 | 19.98±5.58 | 120.60±5.80 | 45.17±8.90 |
| proV1 | 1808.97±533.26 | 26048.07±3254.38 | 8119.52±931.61 |
| proV2 | 1248.25±267.61 | 20226.51±3164.81 | 5557.57±880.61 |
| proW1 | 67.33±7.67 | 1100.01±205.74 | 296.37±28.02 |
| proW2 | 410.52±136.89 | 7934.30±1992.17 | 1963.72±512.42 |
| proX1 | 2708.60±468.16 | 28803.66±6405.98 | 8606.95±1628.96 |
| proX2 | 1739.62±233.27 | 16464.36±3391.55 | 5773.60±938.28 |
| ygaY1 | 28.40±5.41 | 313.55±46.54 | 65.85±7.87 |
| ygaY2 | 26.85±5.16 | 111.52±9.33 | 41.30±8.68 |

**Table S6: Relative transcript levels of amplicons within and flanking the *proVWX* operon in NT331 Δ*stpA* (NT633). Internal control: *hcaT***

| Amplicon | NT633 0.08 M NaCl | NT633 Hyperosmotic shock | NT633 0.3 M NaCl |
|---|---|---|---|
| nrdF1 | 125.97±10.43 | 193.56±14.99 | 150.77±23.88 |
| nrdF2 | 79.80±17.35 | 142.70±24.42 | 116.26±17.29 |
| proU1 | 53.39±10.33 | 295.26±6.41 | 124.22±8.26 |
| proV1 | 5052.77±2083.77 | 63812.32±8172.40 | 23504.72±7658.21 |
| proV2 | 3465.18±1140.43 | 49570.11±8135.08 | 16194.99±5930.20 |
| proW1 | 182.25±3.84 | 2697.16±538.20 | 850.69±237.53 |
| proW2 | 1147.82±517.95 | 19404.52±4834.93 | 5847.17±2784.05 |
| proX1 | 7488.12±2115.74 | 70727.28±16904.09 | 25217.37±9585.28 |
| proX2 | 4800.36±1181.15 | 40439.05±9029.73 | 16724.05±5495.00 |
| ygaY1 | 76.42±7.41 | 768.07±117.33 | 184.66±34.12 |
| ygaY2 | 72.25±6.94 | 273.21±23.27 | 113.45±9.30 |

**Table S7: Relative transcript levels of amplicons within and flanking the *proVWX* operon in NT331 Δ*rnc* (NT632). Internal control: *rpoD***

| Amplicon | NT632 0.08 M NaCl | NT632 Hyperosmotic shock | NT632 0.3 M NaCl |
|---|---|---|---|
| nrdF1 | 83.09±6.94 | 90.11±13.79 | 82.23±7.34 |
| nrdF2 | 69.94±18.13 | 73.93±16.09 | 59.03±7.40 |
| proU1 | 73.48±12.17 | 80.54±21.69 | 57.72±10.85 |
| proV1 | 559.26±96.36 | 8023.76±2095.41 | 4434.21±530.35 |
| proV2 | 444.98±85.59 | 6715.55±1571.80 | 3394.43±544.03 |
| proW1 | 83.09±14.12 | 453.45±83.01 | 239.58±53.78 |
| proW2 | 157.33±19.22 | 1876.76±403.74 | 872.33±175.58 |
| proX1 | 356.34±107.62 | 5511.92±1414.30 | 2818.13±460.38 |
| proX2 | 277.14±63.50 | 4227.19±823.92 | 2319.72±359.24 |
| ygaY1 | 51.17±6.31 | 80.78±7.12 | 59.70±13.03 |
| ygaY2 | 58.37±4.84 | 68.84±9.64 | 57.59±11.98 |

**Table S8: Relative transcript levels of amplicons within and flanking the *proVWX* operon in NT331 Δ*rnc* (NT632). Internal control: *hcaT***

| Amplicon | NT632 0.08 M NaCl | NT632 Hyperosmotic shock | NT632 0.3 M NaCl |
|---|---|---|---|
| nrdF1 | 143.94±18.40 | 151.32±12.66 | 142.88±14.94 |
| nrdF2 | 119.79±24.13 | 123.63±15.83 | 102.38±12.24 |
| proU1 | 126.33±15.92 | 134.26±24.93 | 99.27±9.15 |
| proV1 | 975.93±228.81 | 13700.86±4167.98 | 7699.76±991.49 |
| proV2 | 775.22±183.41 | 11494.71±3369.42 | 5862.75±710.33 |
| proW1 | 143.06±22.42 | 779.19±221.60 | 409.41±48.64 |
| proW2 | 272.42±40.24 | 3200.35±827.86 | 1512.10±312.42 |
| proX1 | 614.23±166.78 | 9329.09±2384.86 | 4897.48±877.88 |
| proX2 | 477.58±95.71 | 7217.89±1818.43 | 4037.97±734.01 |
| ygaY1 | 88.16±8.82 | 137.41±22.07 | 102.15±12.71 |
| ygaY2 | 100.83±10.11 | 116.59±18.27 | 98.49±7.54 |

**Table S9: Relative interaction frequency of proU3_NlaIII with fragments within and flanking the *proVWX* operon in NT331. Cross-linking control: proU3_NlaIII-proU6_NlaIII**

| Interaction fragment | NT331 0.08 M NaCl | NT331 Hyperosmotic shock | NT331 0.3 M NaCl |
|---|---:|---:|---:|
| **proU17_NlaIII** | 38.08±6.78 | 34.77±0.90 | 42.64±2.06 |
| **proU16_NlaIII** | 48.38±9.33 | 40.35±1.19 | 47.88±4.10 |
| **proU13_NlaIII** | 110.91±13.79 | 100.89±2.51 | 122.07±8.30 |
| **proU1_NlaIII** | 72.52±5.73 | 63.02±2.23 | 76.42±2.95 |
| **proU2_NlaIII** | 155.34±17.46 | 123.83±8.94 | 141.17±8.83 |
| **proU4_NlaIII** | 81.39±8.21 | 81.91±3.90 | 88.48±5.82 |
| **proU5_NlaIII** | 140.22±5.60 | 130.55±1.25 | 136.32±4.47 |
| **proU6_NlaIII** | 100.00 | 100.00 | 100.00 |
| **proU7_NlaIII** | 95.91±5.04 | 98.59±8.97 | 100.39±5.33 |
| **proU8_NlaIII** | 35.10±0.43 | 37.75±1.66 | 39.88±1.64 |
| **proU9_NlaIII** | 28.22±1.86 | 27.64±1.48 | 31.33±1.02 |
| **proU10_NlaIII** | 47.43±3.70 | 41.79±3.08 | 47.15±3.09 |
| **proU11_NlaIII** | 51.98±4.69 | 43.40±3.16 | 48.07±1.88 |
| **proU12_NlaIII** | 47.34±3.15 | 45.05±4.16 | 52.05±2.86 |
| **proU14_NlaIII** | 28.11±2.64 | 23.39±1.73 | 28.24±1.13 |
| **proU15_NlaIII** | 32.75±4.00 | 26.70±2.68 | 33.50±2.23 |

**Table S10: Relative interaction frequency of proU3_NlaIII with fragments within and flanking the *proVWX* operon in NT644. Cross-linking control: proU3_NlaIII-proU6_NlaIII**

| Interaction fragment | NT644 0.08 M NaCl | NT644 Hyperosmotic shock | NT644 0.3 M NaCl |
|---|---:|---:|---:|
| **proU17_NlaIII** | 28.37±3.12 | 34.64±4.01 | 38.61±2.76 |
| **proU16_NlaIII** | 39.09±3.45 | 45.97±7.36 | 43.15±2.83 |
| **proU13_NlaIII** | 86.13±8.13 | 103.83±15.69 | 106.67±5.20 |
| **proU1_NlaIII** | 65.71±8.06 | 66.14±4.66 | 71.05±4.12 |
| **proU2_NlaIII** | 127.74±6.05 | 133.70±20.05 | 126.79±12.71 |
| **proU4_NlaIII** | 101.16±4.66 | 108.39±18.94 | 107.01±8.58 |
| **proU5_NlaIII** | 132.61±3.97 | 139.70±6.27 | 136.66±5.42 |
| **proU6_NlaIII** | 100.00 | 100.00 | 100.00 |
| **proU7_NlaIII** | 89.51±4.09 | 100.21±3.76 | 101.13±10.40 |
| **proU8_NlaIII** | 34.05±2.66 | 38.65±3.39 | 38.61±1.17 |
| **proU9_NlaIII** | 27.78±1.36 | 27.27±0.51 | 28.85±2.32 |
| **proU10_NlaIII** | 39.96±1.85 | 43.84±1.30 | 40.87±2.19 |
| **proU11_NlaIII** | 32.25±3.46 | 35.52±1.18 | 35.34±2.77 |
| **proU12_NlaIII** | 39.88±3.71 | 43.69±1.12 | 45.18±2.85 |
| **proU14_NlaIII** | 25.45±1.86 | 23.33±2.09 | 24.59±2.06 |
| **proU15_NlaIII** | 31.44±2.07 | 30.54±2.76 | 30.73±2.91 |

**Table S11: Relative interaction frequency of proU3_NlaIII with fragments within and flanking the *proVWX* operon in NT331 treated with rifampicin. Cross-linking control: proU3_NlaIII-proU6_NlaIII**

| Interaction fragment | NT331 Rif 0.08 M NaCl | NT331 Rif Hyperosmotic shock | NT331 Rif 0.3 M NaCl |
|---|---:|---:|---:|
| **proU17_NlaIII** | 59.23±6.49 | 59.56±2.51 | 51.78±2.11 |
| **proU16_NlaIII** | 71.31±6.79 | 74.98±0.40 | 65.28±2.37 |
| **proU13_NlaIII** | 124.05±4.71 | 120.12±1.60 | 121.87±4.86 |
| **proU1_NlaIII** | 100.04±3.78 | 98.79±2.27 | 98.35±2.44 |
| **proU2_NlaIII** | 134.81±23.76 | 121.26±3.79 | 143.55±9.33 |
| **proU4_NlaIII** | 62.69±3.61 | 64.38±2.87 | 70.21±3.61 |
| **proU5_NlaIII** | 123.92±7.68 | 113.29±2.40 | 125.70±5.62 |
| **proU6_NlaIII** | 96.23±5.09 | 90.39±0.30 | 94.54±1.18 |
| **proU7_NlaIII** | 100.00 | 100.00 | 100.00 |
| **proU8_NlaIII** | 34.91±2.19 | 35.72±3.19 | 33.77±1.54 |
| **proU9_NlaIII** | 35.29±3.12 | 39.63±0.89 | 37.23±1.77 |
| **proU10_NlaIII** | 48.25±2.36 | 49.68±2.11 | 53.36±3.83 |
| **proU11_NlaIII** | 53.53±3.55 | 53.08±1.06 | 54.71±2.44 |
| **proU12_NlaIII** | 46.52±0.74 | 48.09±0.61 | 47.58±1.29 |
| **proU14_NlaIII** | 30.02±2.04 | 32.06±0.60 | 29.61±1.22 |
| **proU15_NlaIII** | 34.95±1.40 | 35.13±0.71 | 31.08±1.21 |
